# Inhibition of integrin alpha V (CD51) reduces inflammation and transition to heart failure following pressure overload

**DOI:** 10.1101/2022.10.10.511547

**Authors:** Clément Delacroix, Alexandra Achab-Ali, Paul Alayrac, Marine Gandon-Renard, Fatou Dramé, David Sassoon, Jean-Sébastien Silvestre, Jean-Sébastien Hulot

## Abstract

**Background:** Integrins are surface receptors that bind to extracellular matrix ligands and regulate cellular function through mechanical stress-initiated signal transduction. Integrin alpha V (or CD51) is implicated in myocardial fibrosis and anti-CD51 therapy improves cardiac function and cardiac fibrotic remodeling following myocardial infarction. However, their contribution in non-ischemic pressure-overload induced heart failure has not been established.

**Methods:** We implanted male C57BL/6J wild-type mice with osmotic minipumps containing a combination of AngII (1.44mg/kg/day) and the α1 adrenergic agonist Phenylephrine (PE)(50mg/kg/day) to induce hypertrophic heart failure. Treatment with AngII alone was used as a model of compensated cardiac hypertrophy. Mice treated with PE or saline were used as controls. Animals were treated with daily intraperitoneal injections of the anti-CD51 molecule cilengitide or vehicle. Cardiac echography, flow cytometry, histological, and protein analyses were used to study the development of fibrosis and cardiac adverse remodeling.

**Results:** Mice treated with the combination of AngII and PE showed maladaptive cardiac hypertrophy associated with a fibrotic remodeling and a rapid transition to heart failure. CD51 protein expression and CD51^+^ cell number were increased in the myocardium of these animals. In contrast, mice treated with AngII alone exhibited compensated cardiac hypertrophy with low levels of fibrosis, no signs of congestive heart failure, and no changes in cardiac CD51 expression as well as CD51^+^ cell number. Anti-CD51 therapy in mice receiving AngII + PE significantly reduced the transition to heart failure and the development of cardiac fibrosis. Anti-CD51 therapy notably reduced the recruitment of monocyte-derived pro-inflammatory CCR2^+^ cardiac macrophages, which also showed a high expression of CD51 at their surface. Macrophages sense matrix stiffness and activate a pro-inflammatory response to stiffer substrates, a response that was blunted by anti-CD51 therapy.

**Conclusion:** Anti-CD51 therapy reduces the transition to heart failure in response to pressure overload and modulates the pro-inflammatory and deleterious action of CD51^+^ myeloid cells. We identified CD51 inhibition as a novel therapeutic strategy for reducing the progression of non-ischemic and pressure-dependent heart failure.

**Clinical perspectives:** What is new?

- We observed a pathologic role of the integrin alpha V in causing a maladaptive response to pressure overload.
- A specific pharmacological inhibition of integrin alpha V reduced the transition to heart failure through modulation of the pro-inflammatory and deleterious action of integrin alpha V^+^ CCR2^+^ cardiac macrophages.

What are the clinical implications?

- This study adds to the growing interest in targeting integrins in cardiac disorders by showing a novel immunomodulatory effect.
- Integrin alpha V inhibition should be considered as a novel therapeutic strategy for reducing non-ischemic and pressure-dependent heart failure.

## Introduction

Integrins are heterodimeric surface receptors that mediate cell adhesion^1^. Most integrins can bind to extracellular matrix components and convey critical information on the changes in the mechanical or biochemical properties of the interstitium^1, 2^. Integrins can regulate cellular function through mechanical stress-initiated signal transduction^2^. Among the integrins, integrin alpha V (ITGAV or CD51) can complex with beta subunits to form receptors that bind to proteins containing the arginine-glycine-aspartic acid (RGD) peptides such as fibronectin or vitronectin. These integrins can sense changes in the extra-cellular matrix (ECM) composition and are notably involved in the fibrotic responses in different organs^3, 4^, including the heart^5, 6^. In the cardiac tissue, integrin alpha V is expressed by different cell sub-types including fibroblasts, fibroblasts-like cells identified by PW1 expression, macrophages, and lymphocytes^6–8^. Recent studies have shown that integrin alpha V is implicated in myocardial fibrosis post-myocardial infarction, notably by promoting TGF-beta activity^9, 10^. Of note, anti-CD51 therapy improves cardiac function and cardiac fibrotic remodeling post-myocardial infarction by reducing TGF-beta activation, and the fibroblasts to myofibroblasts conversion^7^. The importance of integrins in different cardiac disorders is progressively recognized, supporting novel therapeutic approaches such as targeting CD51 in ischemic hearts. However, the contribution of integrins in non-ischemic pressure-overload-induced heart failure has not been clearly established.

In response to pressure overload, the heart develops an adaptive response with an increase in cardiac muscle mass and microvascular density in order to maintain cardiac function^11^. A transition to maladaptive cardiac hypertrophy can also occur and involves the development of cardiac fibrosis, abnormal diastolic function, eventually followed by systolic dysfunction^11, 12^. In addition to cardiomyocyte-specific signaling, the mechanisms driving the compensated vs. decompensated states of cardiac hypertrophy involve a mechano-sensitive activation of fibroblasts and immune cells including monocytes-macrophages^13, 14^. Cre-mediated deletion of integrin alpha V in PDGFRβ^+^ interstitial cells attenuated myocardial fibrosis in a model of angiotensin II infusion^5^. RGD-binding integrins are thus important candidates to act as mechanosensors that regulate the responses to increased mechanical stress.

Here, we characterized the contribution of CD51-expressing cells in different models of chemically-induced pressure-overload leading to adaptive or maladaptive cardiac hypertrophy. We showed, for the first time, that a specific pharmacological inhibition of CD51 abrogated the transition to heart failure through modulation of the pro-inflammatory and deleterious action of CD51^+^ cardiac macrophages. We identified CD51 inhibition as a novel therapeutic strategy for reducing non-ischemic and pressure-dependent heart failure.

## Methods

### Mouse Models

All procedures and animal care were approved by our institutional research committee (CEEA34 and French ministry of research, N° 2019072415398743) and conformed to the animal care guideline in Directive 2010/63/EU European Parliament. All animals received human care in compliance with the “Principles of Laboratory Animal Care” formulated by the National Society for Medical Research and the “Guide for the Care and Use of Laboratory Animals” prepared by the Institute of Laboratory Animal Resources and published by the National Institutes of Health (NIH Publication No. 86-23, revised 1996). C57Bl/6J male mice of 9 to 11 weeks old from Janvier Labs were used. Osmotic pumps were implanted subcutaneously in the anesthetized mouse back via inter-scapula incision. Two main protocols were used in which mice were separated into 4 groups. The first model is based on the delivery of Angiotensin II (1.44mg/kg/day) for 28 days. The second model is based on the combined delivery of Angiotensin II (1.44mg/kg/day) and Phenylephrine (50mg/kg/day). A final group with single Phenylephrine delivery (50mg/kg/day) was used to validate the synergic effect of dual Angiotensin II + Phenylephrine stimulation. Delivery of phosphate buffer saline was used as a control. Mice were then treated with daily IP injection with either RGD-cyclic molecule cilengitide (10mg/kg/day) or vehicle, until sacrifice.

For the study of the PW1 expression, we used Pw1^nLacZ^ mice of C57Bl6/J strain (the gene expression is coupled with a LacZ reporter), crossed with Wild-Type C57Bl/6J as previously reported^8^. The LacZ reporter does not affect the mouse phenotype.

### Cardiac echography

Cardiac echography was performed on anesthetized mice using a Vevo 2100 high-resolution ultrasound device (Visualsonics, Toronto, Canada) with a 40MHz probe (MS-550). Mice were anesthetized with 3% isoflurane in the air for induction and maintained with 1.5%. Mice were depilated in the thoracic region and then placed in a supine position on a dedicated heating platform, allowing monitoring of ECG, temperature, and respiratory frequency. Three Time Motion acquisitions from Parasternal long axis views were recorded. Left ventricle dimensions (internal diameter, septum, and posterior wall thickness) were measured using the VevoLab Software (Visualsonics) that provided associated calculations (Fractional Shortening, Ejection Fraction, Cardiac Output). Mitral, aortic and pulmonary flows were recorded using pulse-wave Doppler mode providing velocities parameters (peak and mean velocities).

### Cell isolation and cytometry

Cells were isolated from the whole left ventricle following enzyme digestion. Briefly, mice were euthanized and the heart was excised. The atrium and the right ventricle were cut out and the left ventricle was rinsed out of its blood. The left ventricle was then cut into small pieces (0.5-1mm^3^) using a scalpel and surgical scissors and directly placed into 2mL of enzymatic solution: Collagenase II – 500U/mL (WOLS04177, Worthington), DNase I – 1,6U/mL (11284932001, Roche), in HBSS with Mg^2+^/Ca^2+^ for 15 minutes at 37°C. The tissue was then quickly homogenized using trituration with 1000μL pipette. The supernatant was extracted and put in STOP solution containing: HBSS with Mg^2+^/Ca^2+^ + 10% FBS (Hyclone FBS (U.S.), GE Healthcare Life Sciences, SH30071.03), on ice. Pellet was filled with a new enzymatic solution for another 15min and the process was repeated until complete digestion. Cell homogenate was then filtered on a 70μm strainer and stained with selected antibodies (listed in supplemental table 1). The different populations were gated, analyzed on a BD LSRFortessa™ X-20 Cell Analyzer.

### Blood pressure assessment

The blood pressure was taken from the animals tail using a photoplethysmograph (BP-2000 system, Visitech). Briefly, the mice are taken to a specific room for pressure measurement, sheltered from noise. After a few minutes of adaptation, the mice are delicately placed inside the individual containment boxes leaving the tail protruding to allow analysis by the photoplethysmograph. The temperature of the station is set at 38°C to keep the mice warm and promote correct dilation of the caudal artery. The animals are left inside the containment boxes for a few minutes prior to the analysis to get used to the environment. Then, the analysis protocol is set as follows:

- 3 adaptation cycles without recording followed by 7 measurement cycles, with a 15 second break between the 4th and 5th measurement cycle.

### Histological analysis

Fibrosis assessment was performed using picrosirius red staining on 4μm FFPE cuts. Images were taken using Hamamatsu scanner at 20X and analysis was performed using both in-house ImageJ macro or FIBER-ML program developed in our center.

Cardiomyocyte size and capillary density were assessed using WGA (FL-1021, Vectorlabs) to tag cardiomyocyte plasma membrane and CD31 (DIA-310, Dianova) to tag endothelial cells. Images were taken using confocal microscopy (Leica SPE) at 40X and analyzed using in-house ImageJ macro.

### Western-Blot

Proteins were extracted from frozen mouse left ventricles using the TissueLyser LT (Qiagen) with 5mm stainless steel beads (69989, Qiagen) into ice-cold RIPA buffer (50 mM Tris pH 7.4, 150 mM sodium chloride, 1% IGEPAL CA-630, 50 mM deoxycholate, and 0.1% SDS) supplemented with anti-proteases (Sigma-Aldrich), anti-phosphatase inhibitors (Sigma-Aldrich), and 1 mM Na3VO4. Samples mechanical digestion was performed with TissuLyser LT set on a shaking frequency of 50Hz, with a cycle of 1min, repeated if needed, until full dissolution of the tissue. After 1 h incubation at 4°C on a spinning wheel, the homogenates were centrifuged at 4°C for 15 min at 15,300 g, and the supernatants containing proteins were collected. Protein concentrations for all samples were determined with Pierce BCA Protein Assay (Thermo-Fisher). Proteins were denatured with NuPAGE LDS sample buffer and NuPAGE Sample Reducing Agent and then loaded on a precast 4-12% Bis-Tris gel (NuPAGE Novex, Life Technologies). After migration at 100 V, proteins were transferred onto nitrocellulose membranes by wet transfer (Bio-Rad). After Ponceau red staining to evaluate transfer’s efficiency, membranes were blocked for 1 h in 5% skim milk in TBS with 0.1% Tween-20 and incubated overnight at 4°C with primary antibodies specific for CD51 (1:1000, # ab179475, Abcam) and Vinculin (1:1000, # V9131, Sigma-Aldrich) used for loading normalization, diluted in 5% milk/TBS-Tween. HRP-conjugated secondary antibodies (1:10000, # 111-035-144, Jackson Immnuo ; 1:10000, # 7076S, Cell Signaling) were used to tag primary antibodies. Immunoreactive bands were detected with an enhanced chemiluminescent procedure using ECL WB Substrate (SuperSignal West Pico Plus, Thermo-Fisher) before imaging with the Amersham ImageQuant800 (Cytiva) and analysis using ImageQuantTM TL software.

### *In vitro* bone marrow-derived macrophage differentiation

Briefly, bone marrow was extracted from 11-week-old C57Bl6/J mice femur and sterilized by a brief dip in ethanol and sterile Phosphate Buffered Saline (PBS, Gibco™) then sectioned in half. Bone marrow was collected by centrifugation (10,000g for 30 sec at 4°C), then cells were filtered using a 70μm strainer and resuspended in growth medium (Gibco™ Medium RPMI 1640, 10% FBS, 1% Penicillin Streptomycin, 5.000 U/mL). To stimulate macrophage differentiation, the growth media was supplemented with recombinant mouse Macrophage Colony Stimulating Factor (M-CSF) at a concentration of 100 ng/mL (416-ML, Bio-techne). Cell density was determined with a cell counter and the cells were seeded into T75 for 7 days at 37°C in 5% CO_2_ at a density of 1×10^6^ cells/cm^2^.

### Differential matrix stiffness macrophages culture

Briefly, after 7 days of differentiation, M0 macrophages were trypsinized and reseeded at a density of 1.2×10^6^ cells/well on P6 wells with either 8 kPa or 16 kPa hydrogel bottom, coated with vitronectin (0,5μg/cm^2^). Macrophages were incubated for 5 hours to adhere to the matrix, then treated with cilengitide (10μM) or vehicle (PBS) for 24 hours. The supernatant was then collected, aliquoted, and snap-frozen for further analysis.

### Immunoflex assay

The supernatant collected from *in vitro* macrophages culture was used undiluted with Bio-Rad Bio-Plex Pro Mouse Cytokine 23-plex Assay (#M60009RDPD) kit, according to the manufacturer protocol. The plate was analyzed on a Bio-Plex 200 system.

### Statistical analysis

The data are expressed as mean ± standard deviation (SD). Pairwise one-way analysis of variance (ANOVA) followed by Tukey’s post-hoc test for multiple comparisons were performed for in three groups comparisons, and Kruskal-Wallis followed by Dunn’s post-hoc test was used if one or more groups did not show normal distribution. The data sets from the HHF model treated with cilengitide vs. vehicle were analyzed using a two-way ANOVA followed by a Tukey post-hoc test. A value of p<0.05 was considered statistically significant. GraphPad Prism 9 software (GraphPad Software, San Diego, CA) was used for the statistical analyses.

## RESULTS

### Models of adaptive vs. maladaptive cardiac hypertrophy

We evaluated the development of cardiac hypertrophy, fibrosis, and dysfunction in different mouse models of chemically-induced pressure overload. Male C57BL/6J mice were implanted with osmotic minipumps containing either the vasoconstrictor Angiotensin II (AngII, 1.44mg/kg/day), a combination of AngII and the α1 adrenergic agonist Phenylephrine (PE, 50mg/kg/day), or phosphate buffer saline as a control (Fig 1A). The duration of administration was planned for 28 days but was reduced to 14 days in the group receiving the dual stimulation because of high mortality and a significant transition to heart failure observed as early as Day 14 (Fig 1A).

**Figure 1:**
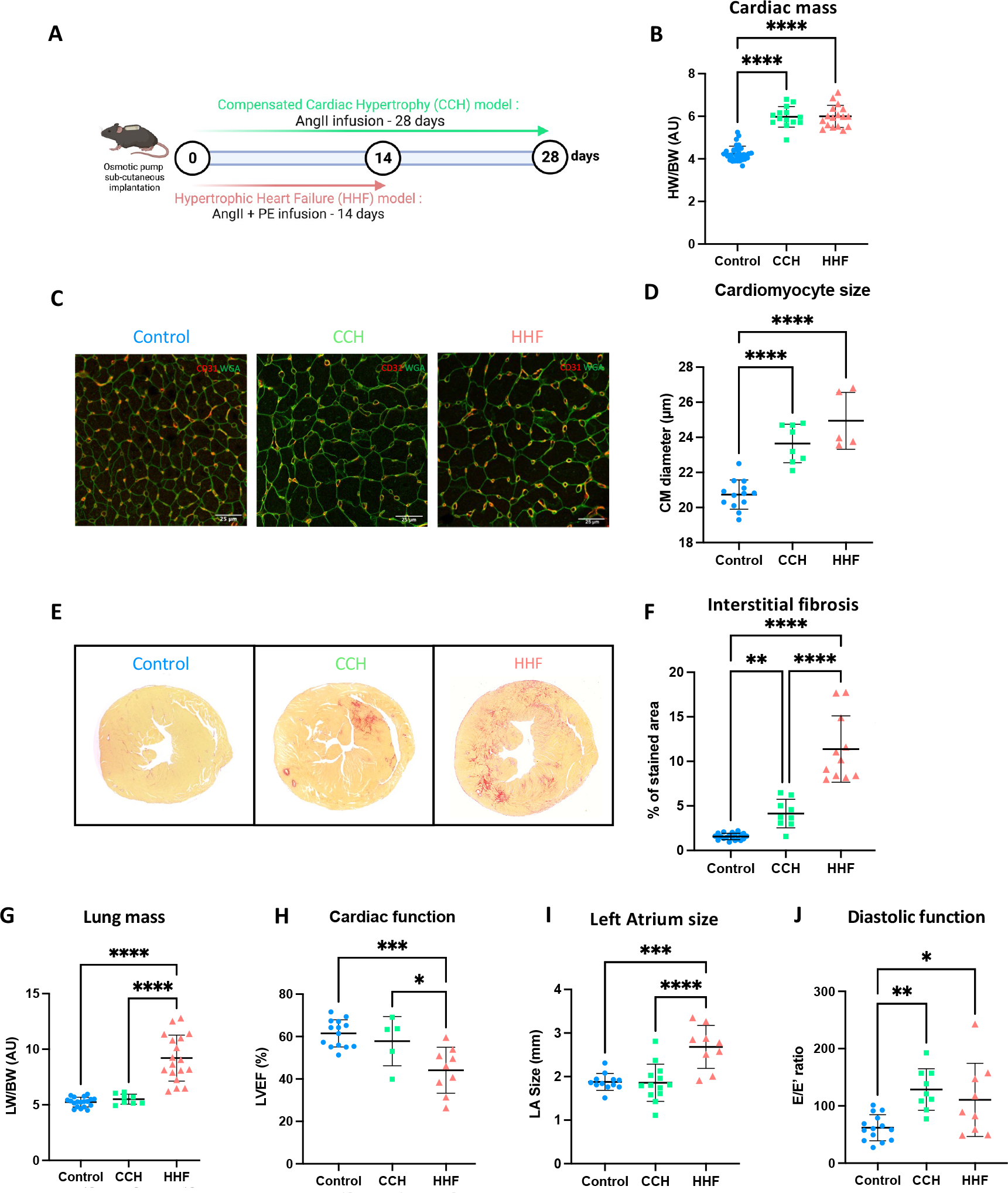
HHF model induces cardiac fibrosis with a transition to heart failure. **A,** Experimental design of pharmacologically-induced models of compensated cardiac hypertrophy (CCH, green) and hypertrophic heart failure (HHF, red). **B,** Cardiac mass in controls (blue), CCH and HHF mice, as measured by heart weight on body weight ratio. N=14 to 35 per group **C,** Immunofluorescent staining of the cardiomyocytes membrane by WGA (green) and endothelial cells by CD31 marking (red) on FFPE cardiac cross-sections and **D,** corresponding measures of cardiomyocyte (CM) diameters. N=5 to 13 per group. **E,** Picrosirius Red staining of collagen fibers on FFPE cardiac cross-sections and **F,** corresponding quantification of the amount of interstitial fibrosis. N=9 to 20 per group **G,** lung weight on body weight ratio. N=8 to 19 per group **H,** Left ventricular ejection fraction (LVEF), **I,** left atria (LA) size, and **J**, E/E’ ratio assessed via cardiac echography. N=5 to 14 per group. Data are presented as Mean ± SD, *p<0.05, **p<0.01, ***p<0.001, ****p<0.0001.

We identified the mouse model of single AngII stimulation as a compensated cardiac hypertrophy (CCH) model and the model with dual stimulation (AngII + PE) as a hypertrophic heart failure (HHF) model. Indeed, we found that the hypertrophic remodeling process was comparable between models either at the organ level, with a similar cardiac mass observed (Fig 1B) or at the cellular level as shown with cardiomyocyte size measurements by WGA staining (Fig 1C-D). However, despite a similar cardiac hypertrophic remodeling and increase in blood pressure (Supp. Fig 1), we found significant differences in the development of interstitial fibrosis between models. Using picrosirius red staining to stain collagen fibers, we found that the HHF model led to a significantly higher level of fibrosis as compared to the levels observed in the CCH model (11.4 ± 3.7% vs. 3.9 ±1.4%, p<0.001, Fig 1E-F). The level of interstitial fibrosis was significantly higher in the CCH model as compared to control, however with levels that stayed in the upper physiological range and remained below 5%. We also found that mice treated with PE alone minorly increased cardiac mass without a significant increase in interstitial fibrosis (Supp. Fig 2A, B), thus identifying a synergic effect of AngII + PE on the development of cardiac fibrosis.

Moreover, on top of the fibrotic remodeling, dual stimulation with AngII + PE was associated with a rapid development of congestive signs in the HHF animals with a significant increase in lung congestion (Fig 1G). Importantly, none of the mice in the CCH model developed significant lung congestion while having similar levels of cardiac hypertrophy (Fig 1B, G). Of note, PE treatment alone improved cardiac mass but did not induce cardiac interstitial fibrosis and lung mass increase (Supp. Fig 2C). The echocardiographic analysis also showed a reduction in systolic function in HHF mice, as well as a significant increase in the left atrium size (Fig 1H-I). Here again, these abnormalities were not found in CCH. However, both CCH and HHF mice had a modified E/E’ ratio (Fig 1J). Capillary density was increased in both CCH and HHF groups (Supp. Fig 3). Together, these results indicate that both CCH and HHF mice display a cardiac hypertrophic remodeling with relaxation trouble, but that the functional consequences and deterioration of congestive heart failure are only observed in the HHF mouse model.

### Increased in CD51 expressing cell number in the failing hearts

We further investigated the mechanisms supporting the differences in fibrosis development and the transition to heart failure between these two models. We have previously shown that cardiac stromal cells expressing integrin alpha V (CD51) are involved in the formation of fibrotic scar in a mouse MI model with permanent LAD ligation^8^. We thus analyzed left ventricle (LV) CD51 protein expression by Western-blot in our cardiac hypertrophy models and found a significant increase in CD51 protein levels in HHF mice when compared to control mice (Fig 2A-B). We also isolated non-cardiomyocyte cells from LV and found a significantly higher number of CD51^+^ cells in the HHF group as seen by flow cytometry analysis (Fig 2C). In line with the limited level of fibrotic remodeling, the number of CD51^+^ cells did not increase in the CCH group. Using different markers (Supp. Tab 1), we then separated non-hematopoietic cardiac cells from hematopoietic cardiac cells in LVs from each group. Interestingly, we found that the number of CD51^+^ cells was enhanced in both non-hematopoietic (Fig. 2D-E) and hematopoietic cardiac cells (Fig. 2F-G) only in HHF mice. These results expand our previous observations post-MI^7^ by revealing an increased amount of CD51 expressing cells in a different experimental model that combines both a fibrotic remodeling and a rapid transition to heart failure.

**Figure 2:**
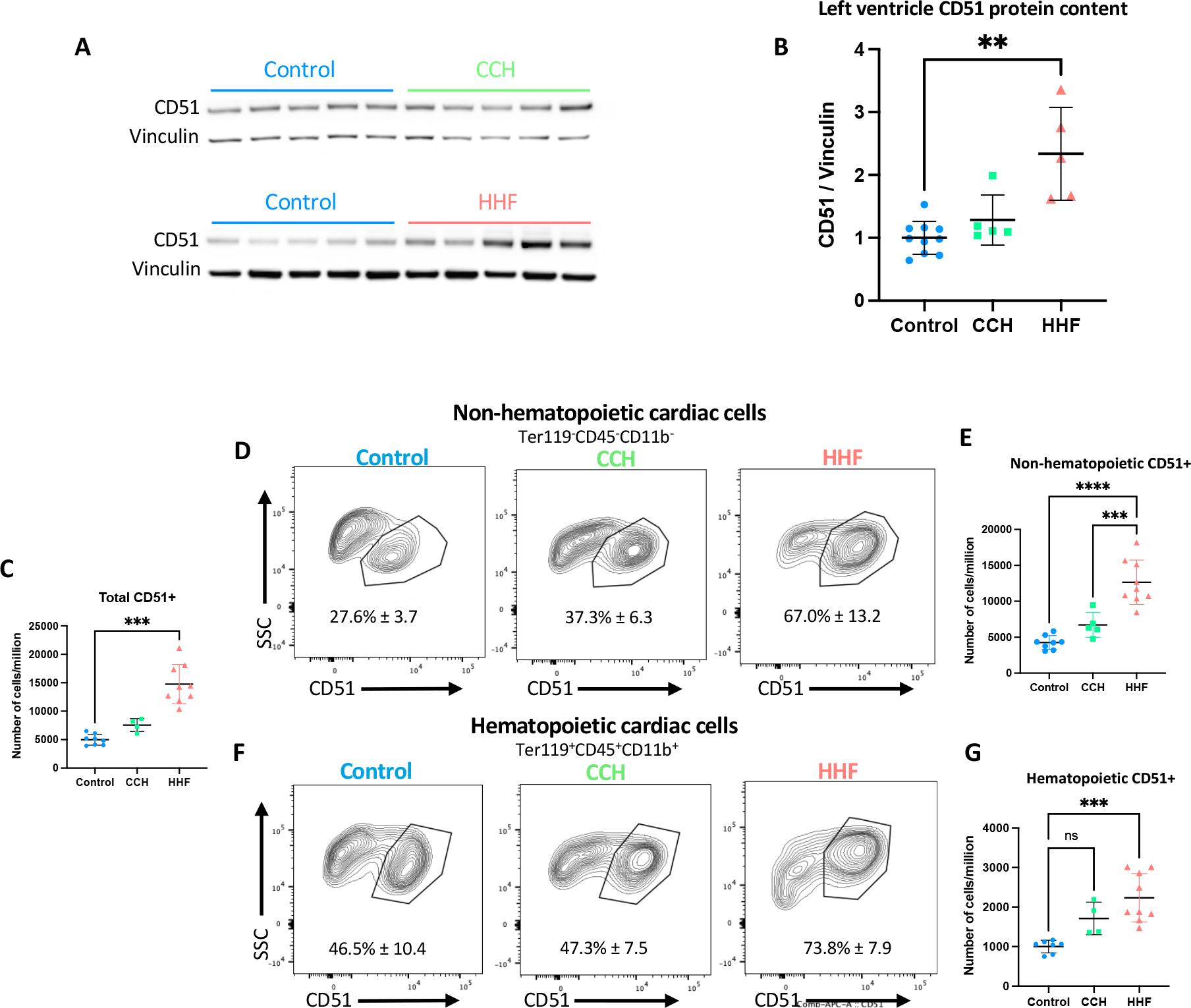
CD51 expression is increased in HHF mice. **A,** Western-Blot analysis of CD51 protein expression in left ventricle of CCH, HHF and control mice, with vinculin as a loading control, and **B,** corresponding quantification. N=5 to 10 per group. **C,** Quantification of total number of CD51^+^ cells found in the myocardium in the different models, assessed by flow cytometry, per million cells analyzed. **D,** CD51 expression in non-hematopoietic cardiac cells, and **E,** corresponding quantification per million cells. **F,** CD51 expression in hematopoietic cardiac cells and **G,** corresponding quantification per million cells. N=4 to 9 per group. **p<0.01, ***p<0.001, ****p<0.0001.

### The anti-integrin CD51 cilengitide reduces fibrosis and prevents the transition to HF

We next tested the preventive effect of CD51 pharmacological inhibition in experimental settings. HHF mice were treated with daily IP administrations of cilengitide or vehicle for 14 days (Fig 3A). We first observed that the cardiac mass was significantly reduced in response to cilengitide at the organ level, as assessed both by gravimetric characterization (Fig. 3B) and echocardiography (Supp. Fig 4A). However, the cardiomyocyte size remained unchanged with the cilengitide treatment (Fig 3C), indicating that this effect is not explained by a simple reduction in the hypertrophic remodeling but rather by a reduction in the content of the interstitial compartment. Cilengitide treatment was associated with a significant reduction in the development of interstitial fibrosis identified by picrosirius red staining (11.4 ± 3.7% vs. 8.6 ±2.8%, p<0.05, Fig 3D-E), without altering perivascular fibrosis (Supp. Fig 5A-B). In addition, cilengitide significantly reduced the transition to heart failure as indicated by a significant reduction in lung mass (Fig 3F) and a preserved LVEF (Fig 3G). Markers of diastolic dysfunction, as well as left atrium enlargement, were unchanged (Fig 3H-I & Supp. Fig 4B).

**Figure 3:**
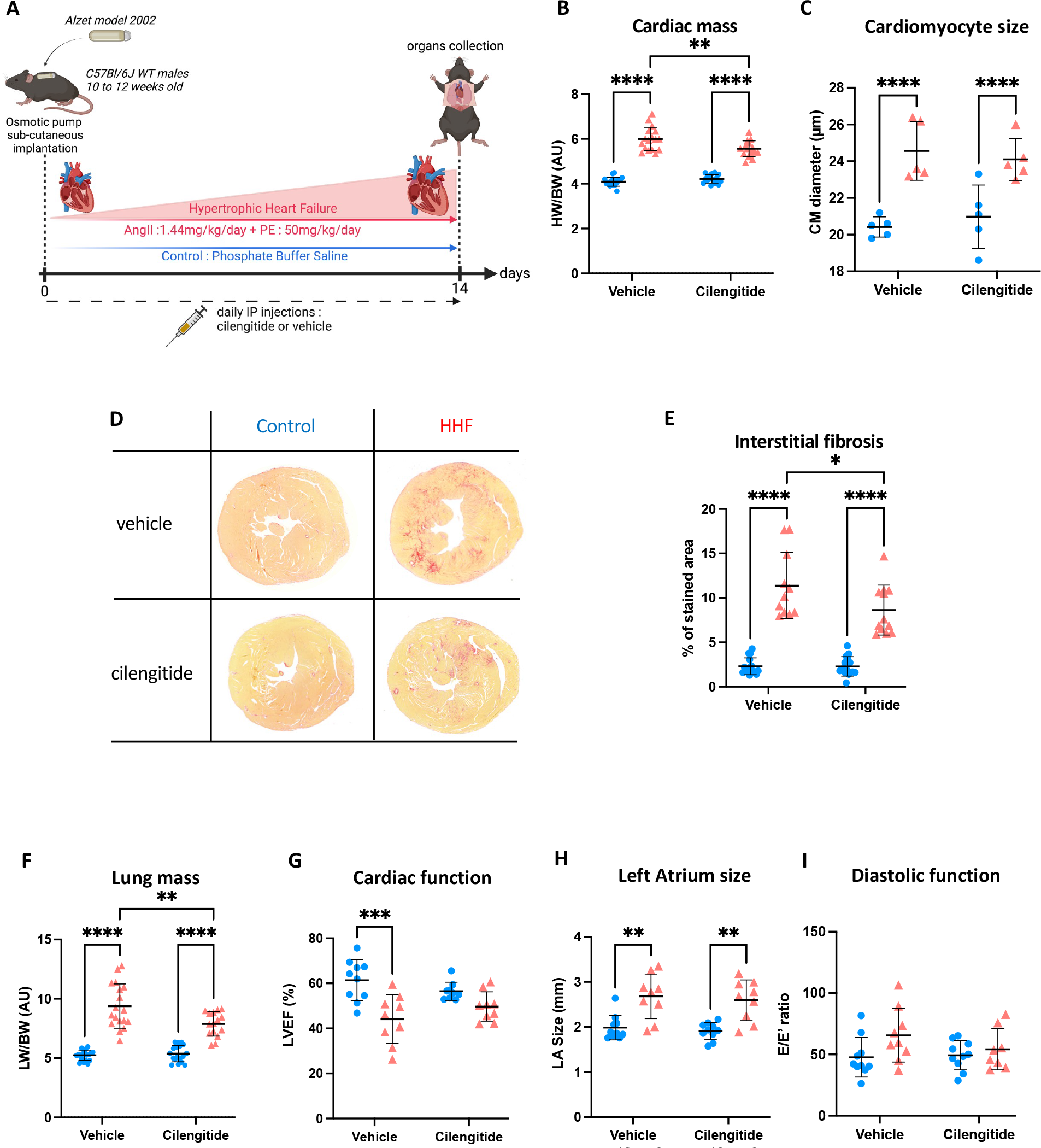
Cilengitide reduces cardiac fibrosis and the transition to heart failure. **A,** Experimental design. Mice were subjected to the HHF (red) or control (blue) protocol and were injected daily with either the CD51 blocker cilengitide or vehicle. **B,** Cardiac mass in the 4 groups of animals, as measured by heart weight on body weight ratio, in control (blue) vs. HHF (red) animals receiving vehicle (left) or cilengitide (right). N=15 to 18 per group. **C,** Quantification of cardiomyocyte size in the 4 groups of animals, measured after immunofluorescent staining of the cardiomyocytes membrane by WGA (green) and endothelial cells by CD31 marking (red) on FFPE cardiac cross-sections. N=5 per group. **D,** Representative images of picrosirius red staining of collagen fibers on FFPE cardiac cross-sections and **E,** corresponding quantification of interstitial fibrosis in the four animal groups. N=12 to 15 per group. **F,** lung weight on body weight ratio in the 4 groups of animals, N=15 to 19 per group. **G,** Left Ventricular Ejection Fraction, **H,** Left Atria size and **I,** E/E’ ratio, assessed via cardiac echography. N=8 to 10 per group. Data are presented as Mean ± SD, *p<0.05, **p<0.01, ***p<0.001, ****p<0.0001

In contrast to the HHF model, cilengitide administered for 28 days by daily IP injection in the CCH model was not associated with any major difference in the development of interstitial fibrosis, cardiac hypertrophy, or cardiomyocyte size (Supp Fig 6A-C). Cilengitide did not change the capillary density or blood pressure in this model (Supp. Fig 6D-E), consistent with a lack of direct vascular impact.

Taken together, these results show that CD51 pharmacological blockade has a preventive impact on the development of both cardiac fibrosis and heart failure development in the context of cardiac hypertrophic remodeling induced by pressure overload.

### CD51 identifies inflammatory monocytes and monocyte-derived macrophages

Concordant with our previous description in the mouse post-MI model^7^, CD51 expression was found in non-hematopoietic cells expressing typical markers of cardiac stromal cells including PDGFR-alpha and PW1 (Supp. Fig 7A-B). The number of those cells increased in the HHF but not in the CCH model (Supp. Fig 7C-D). We also performed a deeper flow cytometry analysis of CD51^+^ cells expressing markers of the hematopoietic compartment (Fig 2F-G) and identified two populations of cells with a high percentage of CD51 expression, namely the monocytes and the macrophages (Fig 4A). Both monocyte and macrophage numbers were increased in HHF but not in CCH mice, confirming the differential involvement of these myeloid cells between these two models (Fig 4B, F). We further investigated the exact nature of the monocytes and macrophages present in the heart of HHF mice to find that a vast majority of the macrophages were CCR2-positive (Fig 4C), thus mostly corresponding to monocyte-derived macrophages with a pro-inflammatory phenotype. Amongst those cells, almost 100% expressed CD51 (Fig 4D-E).

**Figure 4:**
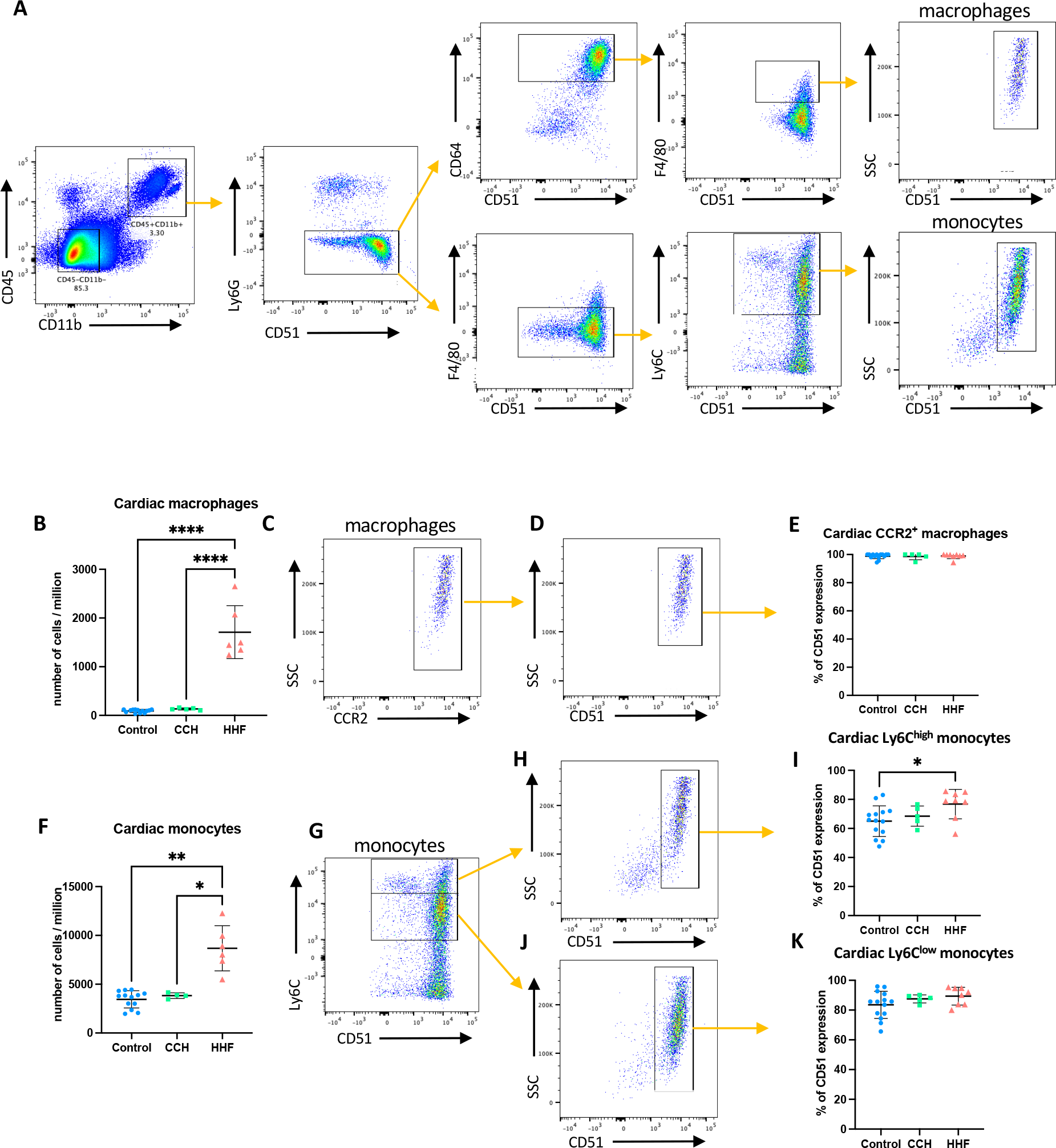
CD51 identifies inflammatory monocyte and monocyte-derived macrophages. **A,** Flow cytometry analysis with sequential narrowing of cells with a high percentage of CD51 expression. **B,** Quantification of F4/80^+^CD64^+^ cardiac macrophages in the CCH, HHF and control models, assessed by flow cytometry, per million cells analyzed. N=5 to 13 per group **C**, Flow cytometry representation of the CCR2 expression in F4/80^+^CD64^+^ macrophages. **D.** Flow cytometry representation of the CD51 expression in the CCR2^+^ macrophage population and **E,** corresponding quantification. N=5 to 14 per group **F,** Quantification of cardiac Ly6C^+^ monocytes in the different models, assessed by flow cytometry, per million cells analyzed. N=4 to 13 per group **G,** Flow cytometry representation of Ly6C^high^ and Ly6C^low^ cardiac monocytes, versus CD51 expression. **H**, Flow cytometry representation of the expression of CD51 in the Ly6C^high^ population and **I,** corresponding quantification. **J**, Flow cytometry representation of the expression of CD51 in the Ly6C^low^ population and **K,** corresponding quantification. N=5 to 14 per group. *p<0.05, a****p<0.0001

Finally, we separated cardiac monocytes according to Ly6C expression levels (low and high) and found that 70% of the Ly6C^high^ monocytes in control and CCH mice and 80% in HHF mice expressed CD51 (Fig 4I), as well as around 90% of Ly6C^low^ monocytes (Fig 4K). Interestingly, the high percentage of CD51^+^ cells was particularly observed in monocytes isolated from the cardiac tissue as compared to those isolated from the spleen, the bone marrow, and the blood (Supp. Fig 8). Together these results indicate that HHF is associated with the increased number of monocytes and macrophages expressing CD51, making them potential targets for anti-integrin therapy such as cilengitide.

### Cilengitide reduces the inflammatory response in HHF animals

Using cytometry analysis, we analyzed changes in the number of CD51^+^ cardiac macrophages and monocytes after cilengitide blockade in HHF mice. Cilengitide significantly reduced the number of CCR2^+^ macrophages as well as Ly6C^low^ monocytes but not that of Ly6C^high^ monocytes (Fig 5A-D). Of note, cilengitide did not modify the amount of Ly6C^+^ monocytes in the blood, the bone marrow, and the spleen, indicating a cardiac-specific effect in the HHF condition (Supp. Fig 9). Altogether, these results indicate that the beneficial effect of the anti-CD51 therapeutic approach could be mediated by the reduction of monocyte/macrophage deleterious effect on cardiac allostasis.

**Figure 5:**
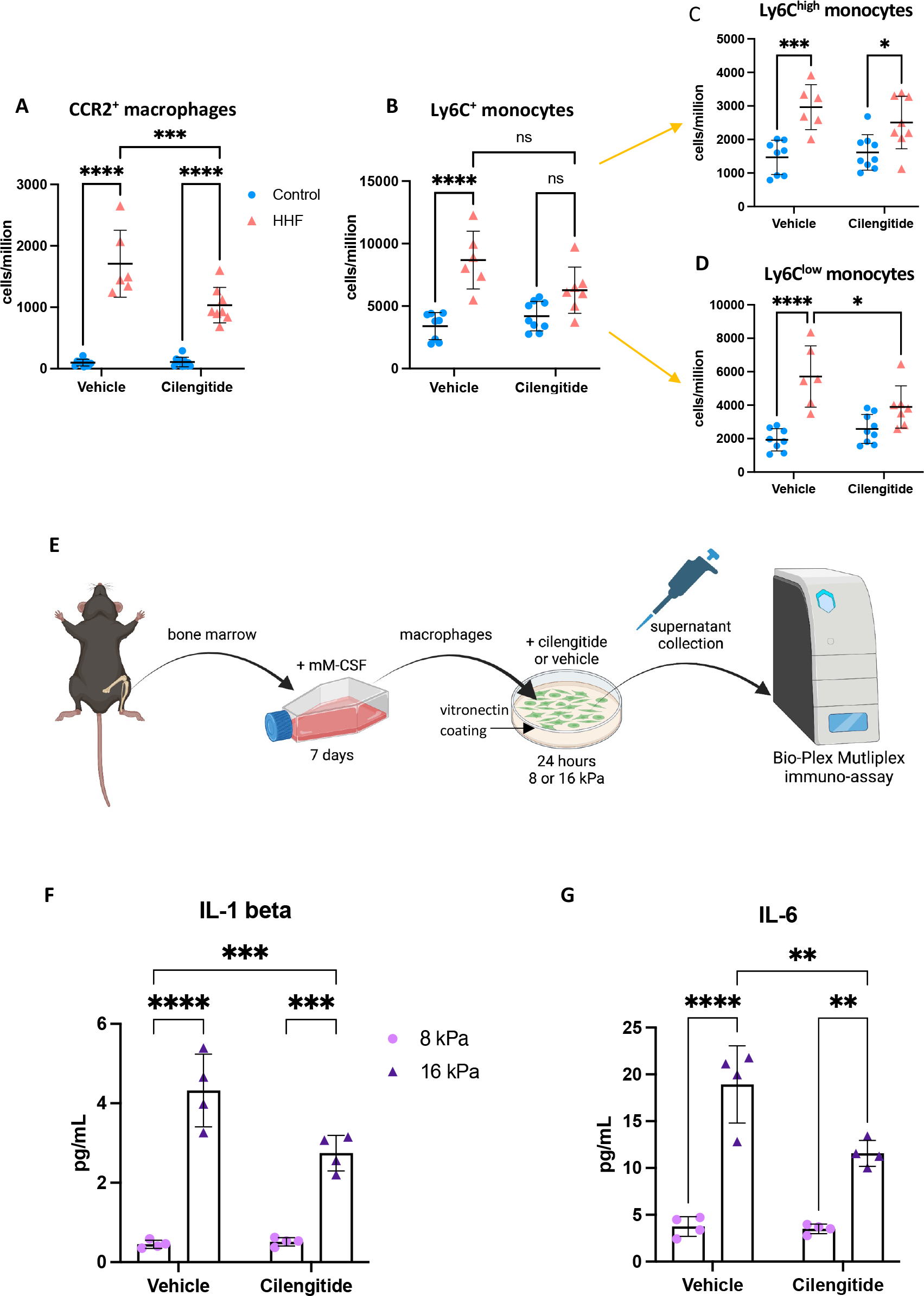
Cilengitide reduces inflammatory cells’ number and pro-inflammatory cytokines’ secretion. **A,** Quantification of cardiac CCR2^+^ pro-inflammatory macrophages, assessed by flow cytometry, per million cells in control (blue) vs. HHF (red) animals receiving vehicle (left) or cilengitide (right). N=6 to 10 per group **B,** Quantification of cardiac Ly6C^+^ monocytes, **C**, of cardiac Ly6C^high^ monocytes, or **D.** cardiac Ly6C^low^ monocytes, assessed by flow cytometry, per million cells N=6 to 10 per group. **E,** Experimental design of the BMDMs culture on different matrix stiffness (8kD vs. 16 kD) using a vitronectin coating and measures of the secreted cytokines ny immunoassay. **F,G** Measures by immunoassay of IL-1 beta (**F**) and of IL-6 (**G**) secreted by BMDMs cultured on soft (8Kd, pink) or stiff (16Kd, purple) vitronectin-coated substrates and treated with vehicle (left) or cilengitide (right); N=4 per group. Mean±SD, *p<0.05, **p<0.01, ***p<0.001, ****p<0.0001

### CD51 blockade shapes macrophage inflammatory phenotype

CD51 is an integrin that mediates adherence to extracellular matrix components, notably vitronectin. We thus interrogated whether CD51 blockade disrupts the interaction of CD51^+^ immune cells with the ECM, and blunts the sensing of a stiffer fibrotic ECM and their consequent cytokine secretion. We differentiated bone marrow cells from 12 weeks old mice for 7 days, into naive M0 macrophages. We then cultivated those macrophages on a vitronectin matrix with two different stiffness (i.e. 8 kPa and 16 kPa), in order to mimic stiffening conditions found in a healthy vs. a fibrotic myocardium, respectively (Fig 5E). Macrophages were then treated for 24 hours with cilengitide or vehicle. The cytokines secretion profile was analyzed in the supernatant using an Immunoflex assay recognizing 23 different analytes (Supp. Tab 2). In line with our hypothesis, we found that the paracrine potential of cultured macrophages was modified in response to a stiffer ECM (Supp Fig 10). Interestingly, we showed that the increase in two prototypic inflammatory cytokines, IL-1 beta and IL-6, was significantly reduced by almost 40% after cilengitide treatment (Fig. 5F-G), suggesting a modulation of the secretion of pro-inflammatory factors by targeting CD51.

## DISCUSSION

Our findings underscore an important role of cells expressing CD51 (i.e., integrin alpha V) in the maladaptive response to cardiac pressure overload. Our results show a significant increase in the expression of CD51, as well as a significant increase in the number of CD51^+^ cells, in animals presenting a rapid transition to HF following chemically-induced pressure overload but not in animals with an adaptive cardiac hypertrophic response. This result was observed despite identical levels of cardiac hypertrophic remodeling in both models, thus suggesting a mechanism independent of cardiomyocyte-specific signaling. We found that CD51 is highly expressed by cardiac stromal cells characterized by typical markers such as PDGFR-alpha or PW1. These cells have been shown to contribute to cardiac fibrosis by favoring myofibroblasts formation^8^ or activating pro-fibrotic pathways related to TGF-beta^7^. Concordantly, a significant increase in the level of cardiac fibrosis was found in the HHF group. However, the absolute levels of interstitial fibrosis were limited (around 10% of total area) and cannot fully explain the rapid transition to HF in these animals, prompting us to investigate other components related to CD51.

Recently, a functional role for monocyte-derived macrophages has been established in the transition from compensated hypertrophy to HF^13, 15^ and we concordantly found a high expression of CD51 in monocytes and macrophages isolated from the cardiac tissues of HHF mice. Notably, the increase in CD51^+^ cells was not observed in the blood, the spleen, or the bone marrow, suggesting a specific increase in the diseased organ. In addition, CD51^+^ macrophages express the C-C Chemokine Receptor type 2. CCR2^+^ macrophages mainly derive from infiltrating monocytes and were recently shown to regulate myocardial inflammation and heart failure pathogenesis^13^. Cardiac macrophages have been implicated in the development of cardiac fibrosis in a CCR2-dependent process^13, 16^.

Previous studies have also reported the expansion of CCR2^+^ monocytes/macrophages during the early phases of cardiac remodeling following pressure overload-induced by a thoracic aortic banding^13, 17^. Concordant with our results, specific and circumscribed inhibition of CCR2^+^ monocytes and macrophages early during pressure overload reduced pathological hypertrophy, fibrosis, and systolic dysfunction during the late phase of pressure overload^13^. Antibody-mediated depletive strategies against CCR2 have been proposed, but a strong limitation of macrophage depletion strategies is the targeting of myeloid populations in other tissues (such as the spleen or the lung) and the related side effects. Similarly, selective CCR2 antagonists can have a systemic impact that will limit its translation to the clinics. Importantly, we found that CD51 was highly expressed by activated cardiac monocytes/macrophages but to a much lower level in other organs, thus suggesting CD51 pharmacological blockade as a more specific way to target CCR2^+^ cardiac macrophages in the hypertrophic hearts. Further experiments are now required to better delineate the therapeutic potential of targeting CD51 in other models of heart failure.

The reduction in heart failure transition in response to a pharmacological blockade of CD51 is another important finding of our study. We have previously shown that cilengitide significantly ameliorates cardiac outcome after MI in a murine model, through the targeting of cardiac stromal CD51^+^ cells and reduced their differentiation in myofibroblasts^7^. Our present results show that the development of cardiac fibrosis was also decreased in our HHF model, thus strengthening evidence that targeting integrins can represent an appealing anti-fibrotic strategy^3, 18^. Similarly, integrin alpha V blockade has been associated with reduced renal, liver, and pulmonary fibrosis^19–21^. Our results now show that CD51 pharmacological blockade could also be used to limit the transition from compensated to decompensated cardiac hypertrophy, which is a novel finding. Here, we propose that a part of this preventive effect relates to a modulation of the pro-inflammatory and deleterious action of CD51^+^ macrophages. It has been suggested that matrix stiffening can regulate macrophage polarization and exacerbates their pro-inflammatory responses^22, 23^, but whether this process is observed in the cardiac tissue is under-explored. Recent reports have shown the presence of mechanically-gated sensors in immune cells, their ability to transduce a mechanical stimulus to these cells, and the consequent activation of a pro-inflammatory response^24^. Concordantly, our results show that macrophages display a pro-inflammatory secretion profile in response to a stiffer substrate (i.e. 16 kPa) that was selected to mimic the mechanical conditions of myocardial tissue with fibrotic remodeling. Our findings show that the sensing of changes in the mechanical constraints is partly conveyed by integrins that are typically known to mediate adherence to extracellular matrix components^2^. Our results add to the growing interest in targeting integrins in cardiac disorders by showing a novel immunomodulation impact. Inflammation is a hallmark of heart failure^25^. However, therapeutic immunomodulation in HF requires a better understanding of the respective role of the different immune cells.

In conclusion, we report on the critical role of CD51-expressing cells in the process of maladaptive cardiac remodeling in response to pressure overload. CD51 expression identifies different cell subsets, including CCR2^+^ monocytes/macrophages. Pharmacological inhibition of CD51 reduces the transition to heart failure and modulates the pro-inflammatory and deleterious action of CD51^+^ myeloid cells. We identified CD51 inhibition as a novel therapeutic strategy for reducing the progression of non-ischemic and pressure-dependent heart failure.

## Author contributions

CD and JSH designed the study with the help of DS and JSS. CD and JSH wrote the manuscript. CD, AA, PA, MGR, and FD performed the experiments. CD and JSH analyzed the data with the help of JSS. DS and JSS provided critical appraisal of the manuscript. JSH obtained funding for the study and supervised the personnel conducting experimentation and data analysis

## Acknowledgments

We thank the Flow cytometry platform and the histology facility at the Paris Cardiovascular Research Center (PARCC) for their assistance. Cardiac echographies were performed at the PIV Université de Paris Cité, Institut Cochin.

## Sources of funding

This work was supported by a grant from ANR PACIFIC (ANR-18-CE14-0032-02), the Fondation Leducq (13CVD01), and a fellowship from Société Française de Cardiologie – GRRC to CD.

Outside the submitted work, JSH is supported by AP-HP, INSERM and is coordinating a French PIA Project (2018-PSPC-07, PACIFIC-preserved, BPIFrance) and a University Research Federation against heart failure (FHU2019, PREVENT_Heart Failure).

## Disclosures

JSH reports research grants to the institution from Bioserenity, Pliant Thx, Sanofi, Servier; speaker, advisory board, or consultancy fees from Alnylam, Amgen, Astra Zeneca, Bayer, Bioserenity, Boerhinger Ingelheim, MSD, Novartis, Novo Nordisk, Vifor Pharma, all unrelated to the present work. Other authors declare no competing financial interests.

## Supplemental Figures

**Supplemental Figure 1: Blood pressure is increased in CCH and HHF mice**

**Supplemental Figure 2: Single PhenylEphrine stimulation does not induce adverse remodeling**

**Supplemental Figure 3: Capillary density is increased in CCH and HHF mice**

**Supplemental Figure 4: Capillary density is increased in CCH and HHF mice**

**Supplemental Figure 5: Perivascular fibrosis is unchanged after cilengitide treatment**

**Supplemental Figure 6: Cilengitide treatment does not ameliorate CCH**

**Supplemental Figure 7: CD51 expressing PDGFR-alpha and PW1 stromal cells number is increased in HHF mice**

**Supplemental Figure 8: CD51 is differently expressed between hematopoietic progenitors**

**Supplemental Figure 9: Monocyte quantity is unchanged by cilengitide in hematopoietic progenitors**

**Supplemental Figure 10: Modulation of cytokines after cilengitide treatment**

**Supplemental Table 1: Flow Cytometry antibody panel**

**Supplemental Table 2: Bio-Rad Bio-Plex Immunoplex 23-plex assay’s analytes panel**

