## Supplemental figures and tables for "Inhibition of integrin alpha V (CD51) reduces inflammation and transition to heart failure following pressure overload"

### Supplementals

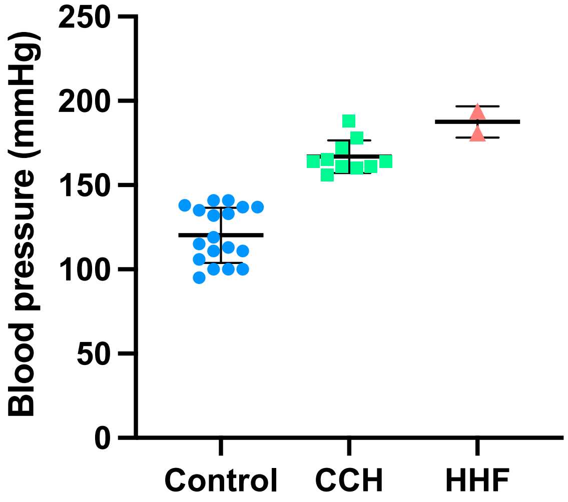


**Supplemental Figure 1:** **Blood pressure is increased in CCH and HHF mice**
Systolic blood pressure assessed at 14 days (CCH) and 7 days (HHF). N=2 to 18 per group.

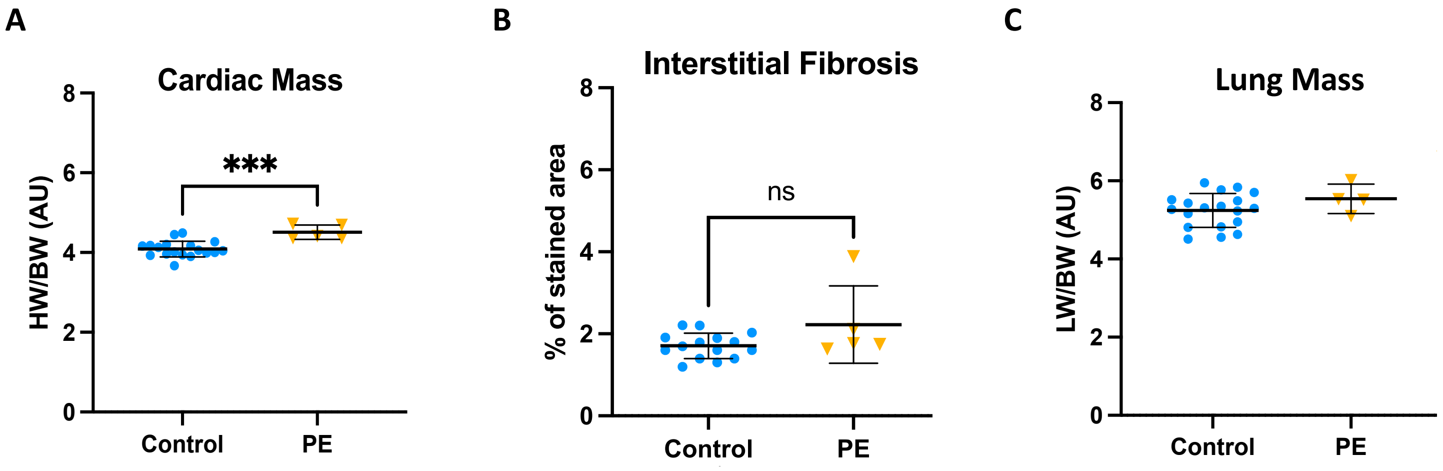


**Supplemental Figure 2 : Single PhenylEphrine stimulation does not induce adverse remodeling**
Measurement of **A,** cardiac mass, assessed by heart weight on body weight ratio. **B,** percentage of interstitial fibrosis measured by picrosirius red staining. **C,** Lung mass assessed by Lung Weight on Body Weight ratio. N=4 to 18 per group. Mean±SD, ***p<0.001.

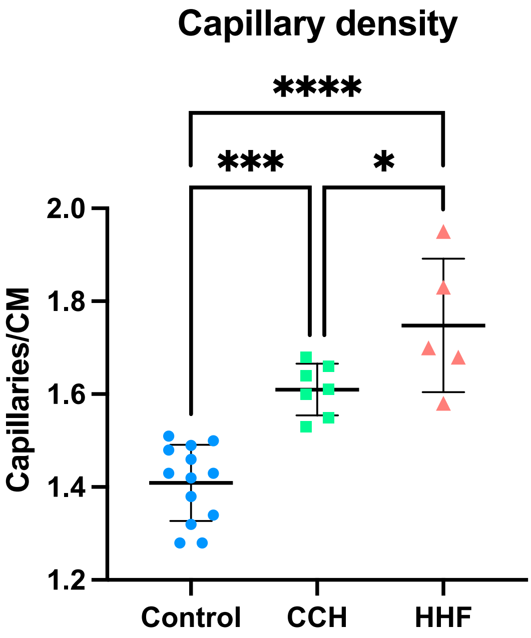


**Supplemental Figure 3 : Capillary density is increased in CCH and HHF mice**
Measurement of capillary density assessed by capillaries on cardiomyocytes ratio. N=7 to 13 per group. Mean±SD, *p<0.05, ***p<0.001, ****p<0.0001.

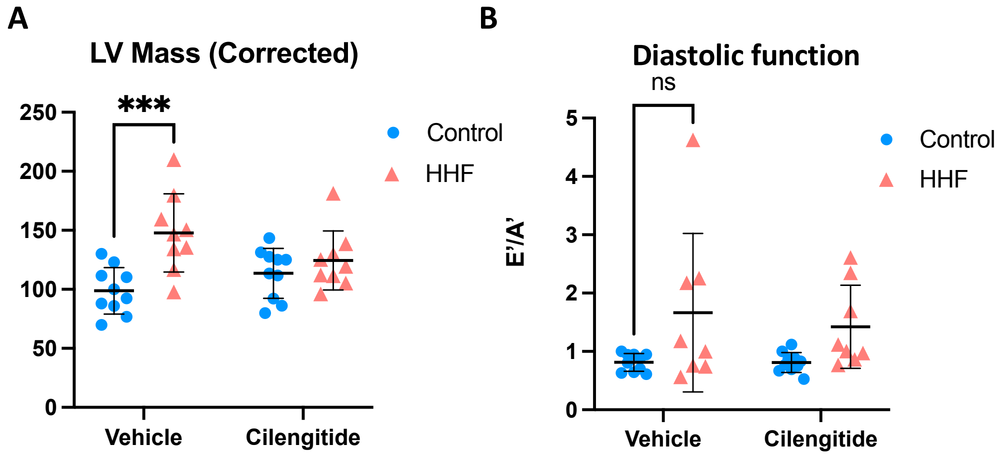


**Supplemental Figure 4 : LV mass increase assessed in cardiac echography is blunted by cilengitide**
**A,** Measurement of the LV mass, as measured by cardiac echography. **B,** Analysis of the E'/A' ratio, as measured by cardiac echography. N=8 to 10 per group. Mean±SD, ***p<0.001, NS = Non-significant.

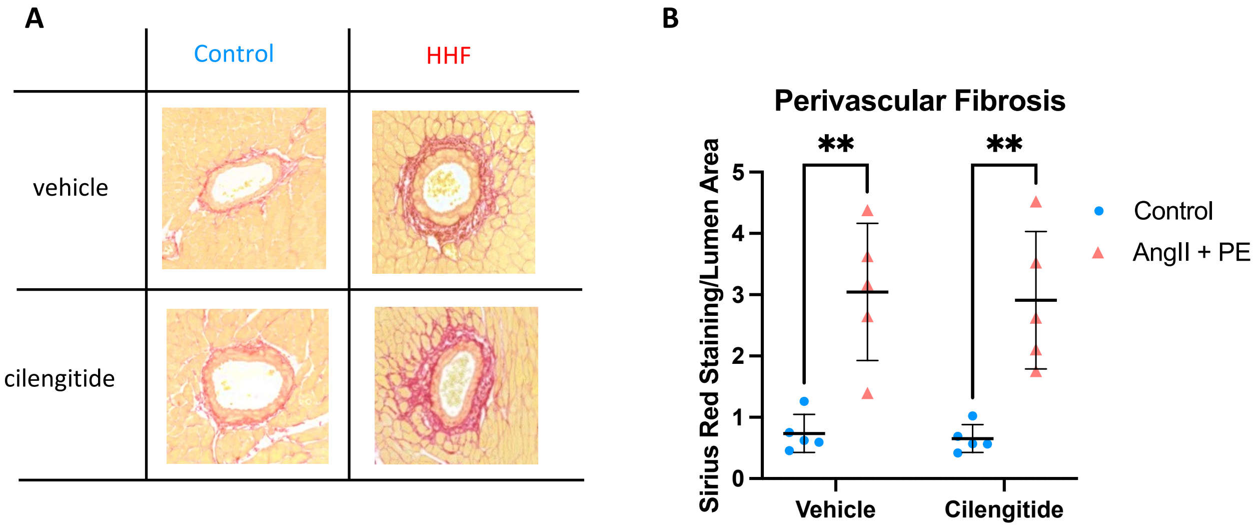


**Supplemental Figure 5 : Perivascular fibrosis is unchanged after cilengitide treatment**

**A,** Picrosirius Red Staining of collagen fibers on FFPE cardiac cross- sections and **B,** corresponding analysis of perivascular fibrosis. N=5 per group. Mean±SD, **p<0.01.

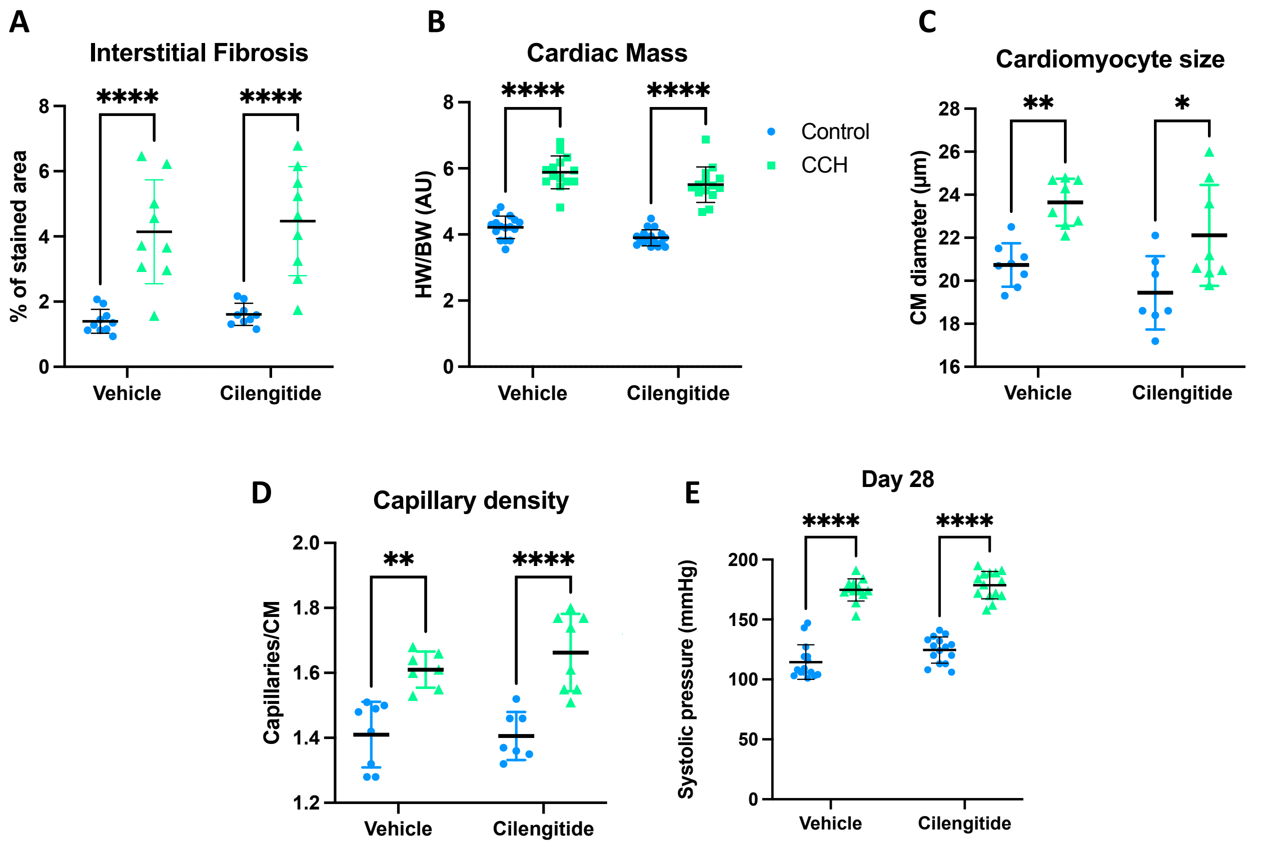


**Supplemental Figure 6 : Cilengitide treatment does not ameliorates CCH**

Measurement of **A,** cardiac mass, assessed by heart weight on body weight ratio. **B,** interstitial fibrosis assessed by picrosirius red staining, **C,** cardiomyocyte size assessed by WGA staining, **D,** capillary density assessed by WGA and CD31 staining and **E,** systolic blood pressure at 28 days (endline) assessed by photoplethysmography. N=7 to 17 per group. Mean±SD, $*p<0.05$, $**p<0.01$, $****p<0.0001$.

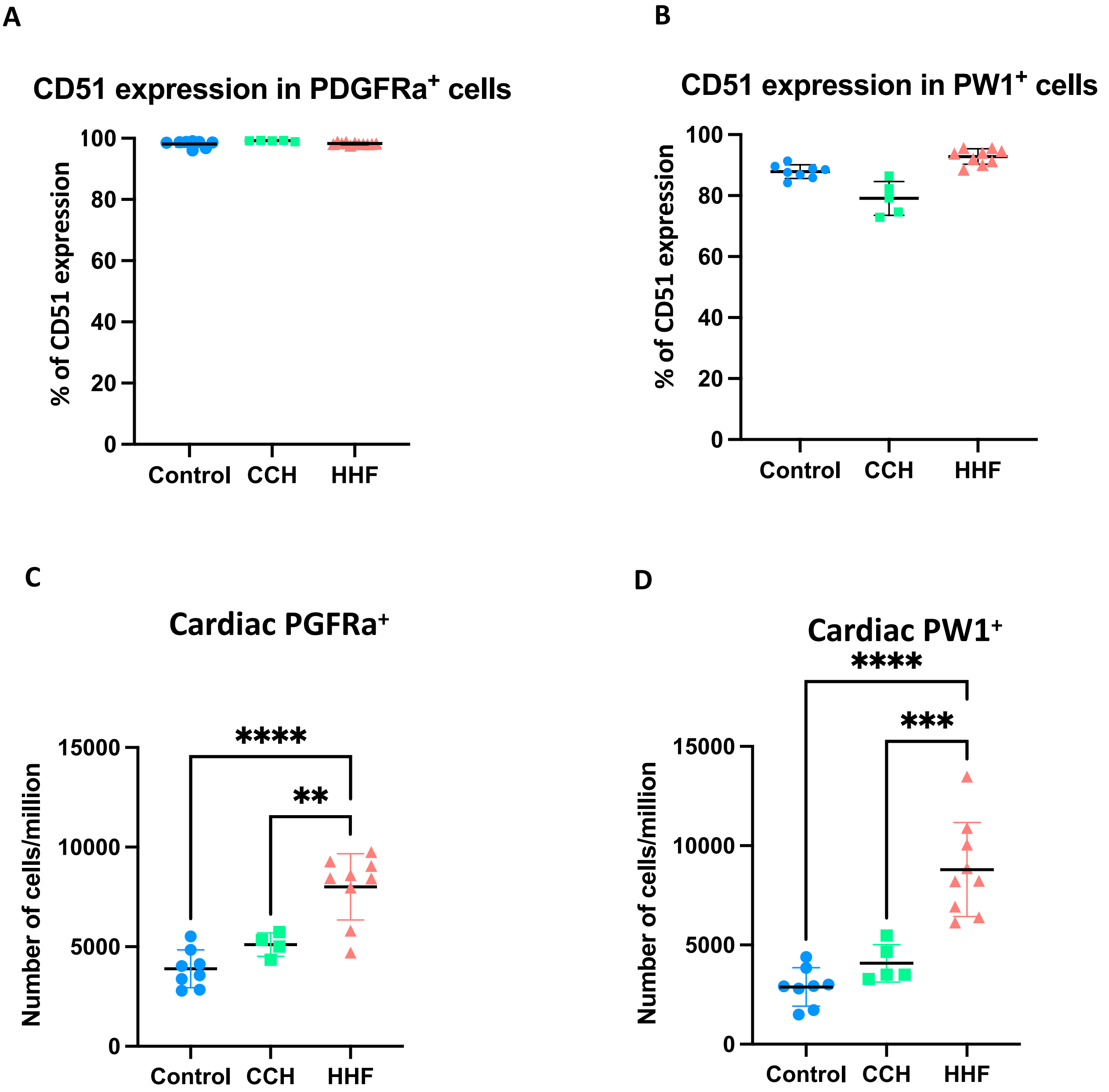


**Supplemental Figure 7 : CD51 expressing PDGFR-alpha and PW1 stromal cells number is increased in HHF mice**
Flow cytometry analysis of the percentage of CD51 in **A,** PDGFRa$^{+}$ cells and, **B,** PW1$^{+}$ cells. Flow cytometry analysis of the quantity of **C,** PDGFRa$^{+}$ cells and **D,** PW1$^{+}$ cells. N=4 to 9 per groups. Mean±SD, **$p<0.01$, ***$p<0.001$, ****$p<0.0001$.

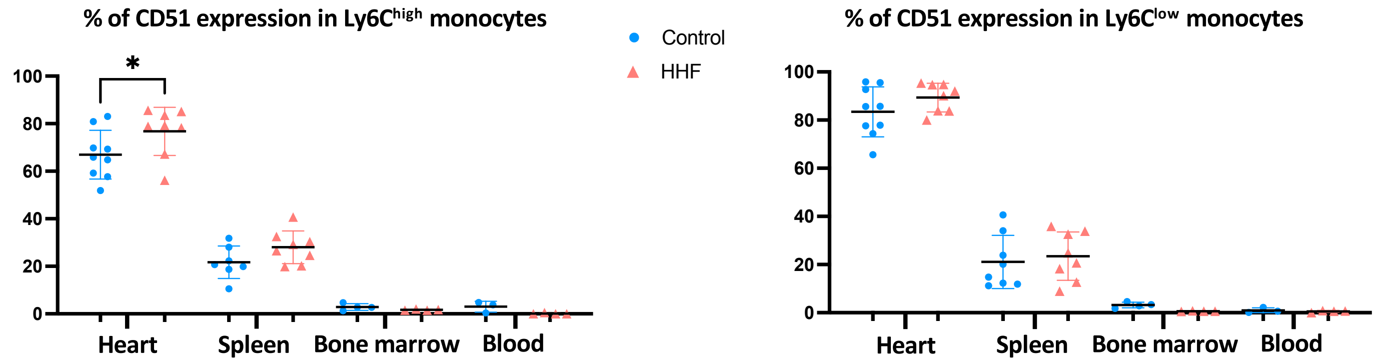


**Supplemental Figure 8 : CD51 is differently expressed in monocytes in different organs**
Flow cytometry analysis of the percentage of CD51 expression in **A,** Ly6C$^{high}$ monocytes and **B,** Ly6C$^{low}$ monocytes in the heart, spleen, bone marrow and blood. N=4 to 9 per group. Mean±SD, *$p<0.05$.

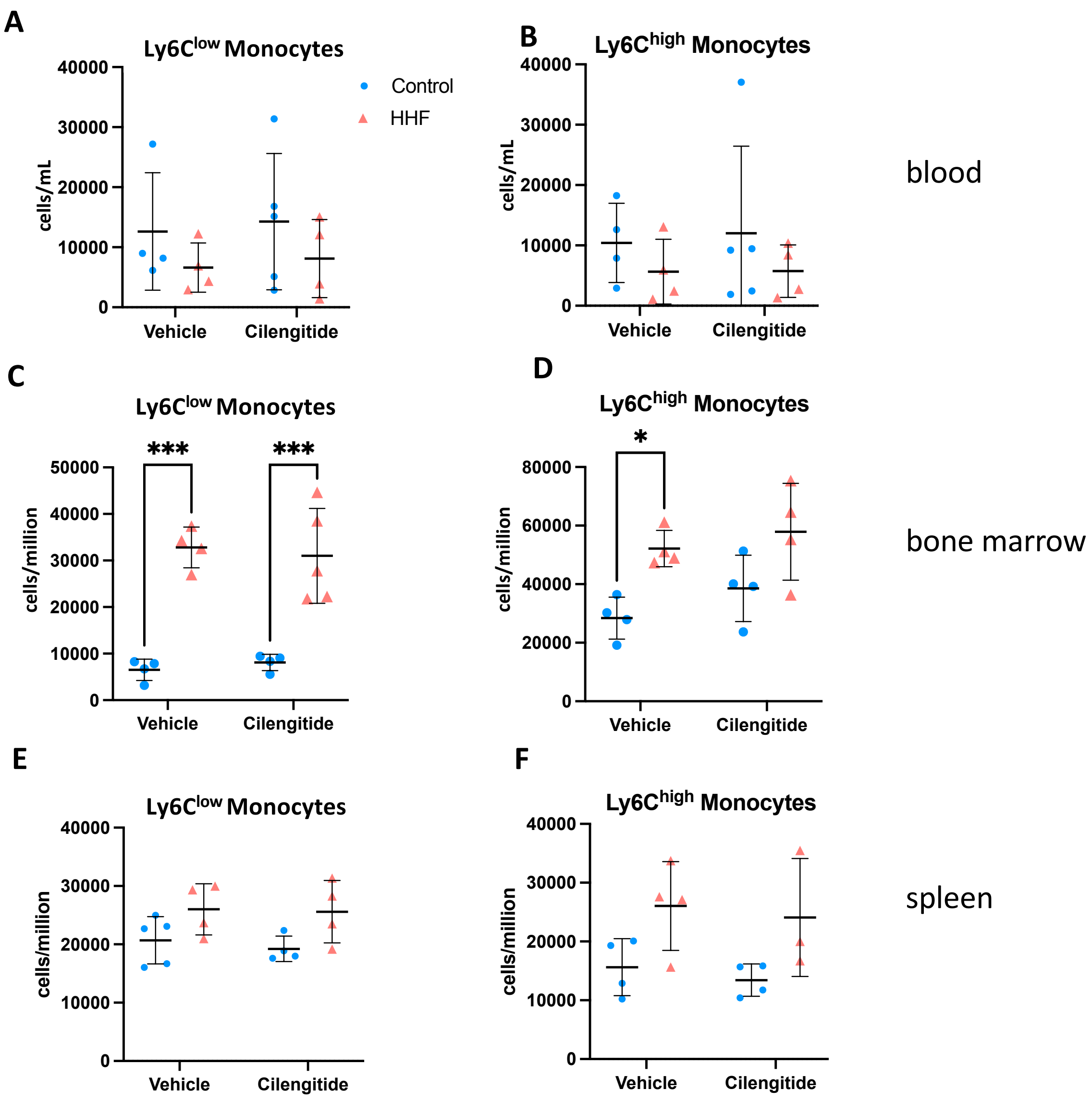


**Supplemental Figure 9 : Monocyte quantity is unchanged by cilengitide**

Flow cytometry assessment of the quantity of **A,** Ly6C$^{low}$ monocytes and **B,** Ly6C$^{high}$ monocytes in the blood, per mL of blood, **C,** Ly6C$^{low}$ monocytes and **D,** Ly6C$^{high}$ monocytes in the bone marrow and **E,** Ly6C$^{low}$ monocytes and **F,** Ly6C$^{high}$ monocytes in the spleen, per million cells analyzed. N=4 to 9 per group. Mean±SD, *p<0.05, ***p<0.001.

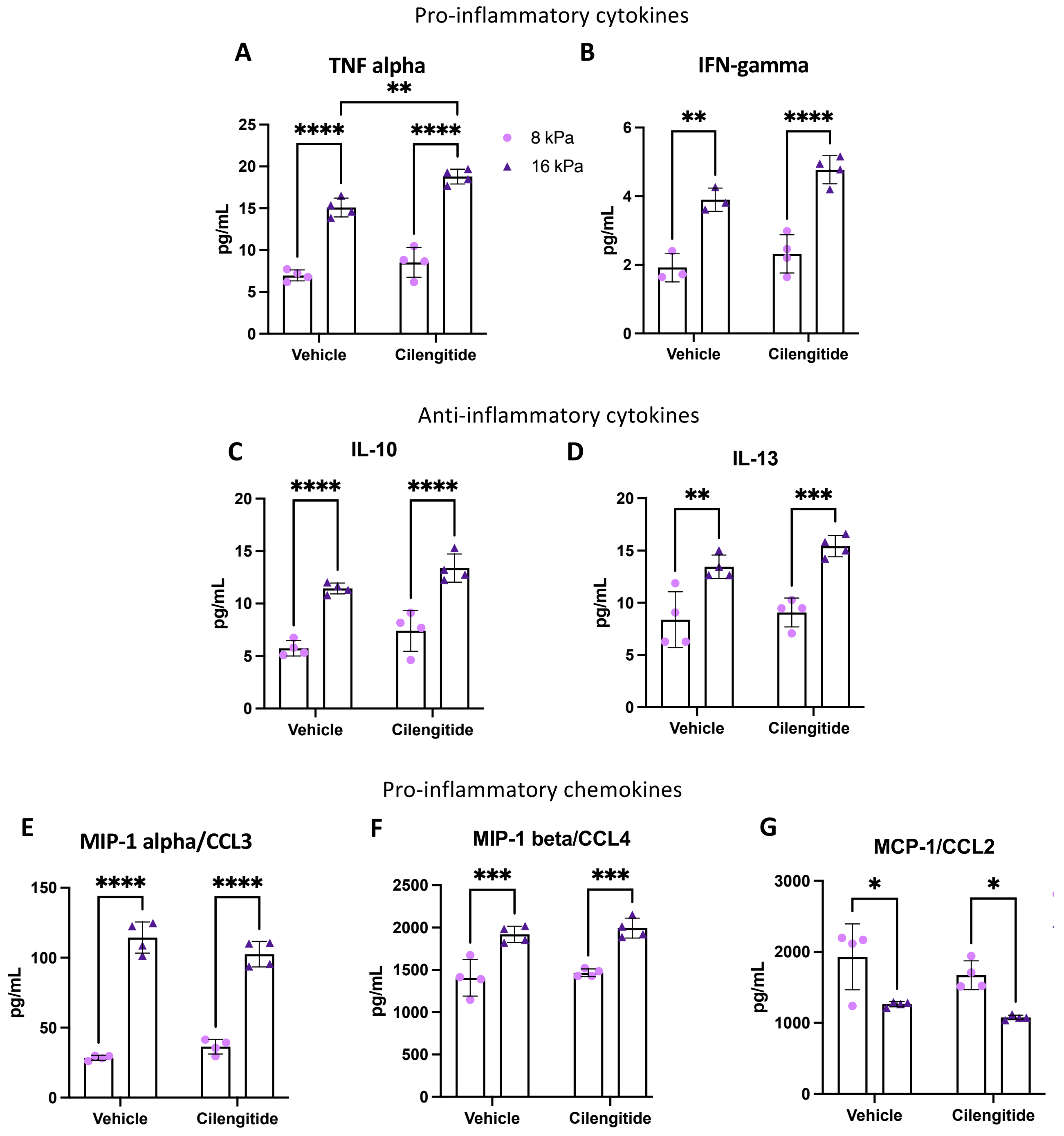


**Supplemental Figure 10 : Modulation of cytokines after cilengitide treatment**
Immunoassay dosage of BMDMs' secretion of **A,** Tumor Necrosis Factor alpha, **B,** Interferon gamma, **C,** Interleukin 10, **D,** Interleukin 13, **E,** Macrophage Inflammatory Protein alpha, **F,** Macrophage Inflammatory Protein beta and **G,** Monocyte Chemoattractant Protein 1, measured by immunoassay. N=4 per group. Mean±SD, *p<0.05, **p<0.01, ***p<0.001, ****p<0.0001.

| Target | Fluorochrome | Clone | Supplier | Dilution |
|---|---|---|---|---|
| Ter119 | BUV395 | TER-119 | BD Biosciences | 1/50[e] |
| CD45 | BUV395 | 30-F11 | BD Biosciences | 1/50[e] |
| CD11b | BUV395 | M1/70 | BD Biosciences | 1/100[e] |
| PDGFR-alpha | PE | APA5 | BD Biosciences | 1/50[e] |
| PW1 (B-gal) | FITC ($C_{12}$ FDG) | / | Invitrogen | 1/300[e] |
| Streptavidin | APC-Cy7 | / | BioLegend | 1/500[e] |
| CD51 Biotin | / | RMV-7 | eBioscience | 1/100[e] |
| CD45 | BV510 | 30-F11 | BioLegend | 1/100[e] |
| CD11b | BV605 | M1/70 | BD Biosciences | 1/100[e] |
| Ly6G | BUV395 | 1A8 | BD Biosciences | 1/100[e] |
| Ly6C | PE | AL21 | BD Biosciences | 1/100[e] |
| CD64 | BV421 | X54-5/7.1 | BioLegend | 1/100[e] |
| F4/80 | AF700 | CI:A3-1 | Bio-Rad | 1/100[e] |
| CCR2 | FITC | SA203G11 | BioLegend | 1/100[e] |
| CD3 | PE-Cy7 | 145-2C11 | BD Biosciences | 1/100[e] |
| CD4 | BV650 | RM4-4 | BD Biosciences | 1/100[e] |
| CD8 | PerCP-Cy5.5 | 53-6.7BD | BD Biosciences | 1/100[e] |
| CD25 | APC | PC61.5 | eBioscience | 1/100[e] |
| B220 | BUV496 | RA3-6B2 | BD Biosciences | 1/100[e] |

**Supplemental Tab 1 : Flow Cytometry antibody panel**

| Cytokines | Chemokines | Growth Factors |
|---|---|---|
| IL-1α | MCP-1/CCL2 | GM-CSF |
| IL-1β | MIP-1α/CCL3 | G-CSF |
| IL-2 | MIP-1β/CCL4 | |
| IL-3 | RANTES/CCL5 | |
| IL-4 | Eotaxin/CCL11 | |
| IL-5 | KC/CXCL1 | |
| IL-6 | | |
| IL-9 | | |
| IL-10 | | |
| IL-12 (p40) | | |
| IL-12 (p70) | | |
| IL-13 | | |
| IL-17A | | |
| IFN-γ | | |
| TNF-α | | |

**Supplemental Tab 2 : Bio-Rad Bio-Plex Immunoplex 23-plex assay's analytes panel**
